# Environmental complexity favors the evolution of learning

**DOI:** 10.1101/019968

**Authors:** Slimane Dridi, Laurent Lehmann

## Abstract

Learning is a fundamental biological adaptation that is widespread throughout the animal kingdom. According to previous research, two conditions are necessary for learning to be adaptive: between-generation environmental variability and within-generation environmental predictability. In this paper, we show that between-generation variability is not necessary, and that instrumental learning can provide a selective advantage in complex environments, where an individual is exposed to a large number of different challenges during its lifespan. We construct an evolutionary model where individuals have a memory with limited storage capacity, and an evolving trait determines the fraction of that memory that should be allocated to innate responses to the environment versus learning these responses. The evolutionarily stable level of learning depends critically on the environmental process, but generally increases with environmental complexity. Overall, our work sheds light on the importance of global structural properties of the environment in shaping the evolution of learning.

## Introduction

Learning allows an individual to use experience and thereby express payoff-relevant actions in novel environments. In particular, instrumental learning permits to build simple associations between newly encountered stimuli and appropriate actions (Pearce, 2008). It is important to understand the adaptive value of this form of learning as it permeates the animal kingdom and underlies the ecological success of the hominin lineage (Johnston, 1982; Boyd and Richerson, 1988; Shettleworth, 2009; Fawcett et al., 2013). In previous work, the selective advantage of learning has been proposed to crucially rely on between-generation environmental variability (Boyd and Richerson, 1988; Stephens, 1991; Feldman et al., 1996; Kerr and Feldman, 2003; Wakano et al., 2004; Dunlap and Stephens, 2009). The argument is that if offspring live in environments where the consequences of actions are totally different from that of parents, and were never experienced in the history of the population, then offspring can express novel appropriate actions only through learning. It has also been emphasized that the environment should not change too fast within an individual’s lifespan for learning to evolve (Stephens, 1991; Dunlap and Stephens, 2009); in other words the environment should be predictable enough for information to be useful.

The general consensus in the literature on the evolution of learning is thus that two conditions are necessary for learning to be adaptive: between-generation environmental variability and within-generation predictability (Boyd and Richerson, 1988; Stephens, 1991; Feldman et al., 1996; Kerr and Feldman, 2003; Wakano et al., 2004; Dunlap and Stephens, 2009). The requirement of predictability seems unavoidable, because learning can be effective only if there is a certain amount of temporal autocorrelation (Fawcett et al., 2014), i.e. if information is reliable over time. The importance of between-generation variability is less clear, however, and we will show in this paper that the occurrence of such variability is not a necessary condition for learning to evolve.

Though learning has indeed been shown to evolve under between-generation environmental variability (e.g. the infinite-environmental state model of Feldman et al., 1996; Wakano et al., 2004), the conditions under which general phenotypic plasticity evolve are very similar (i.e. variable environments, Gomulkiewicz and Kirkpatrick, 1992; Pigliucci, 2001). This blurs the specific advantages of learning over other forms of behavioral plasticity, such as innate behavioral plasticity (Mery and Burns, 2010; Hollis and Guillette, 2011; Snell-Rood, 2013), which is exemplified by fearful reactions to predators, or preference for tasty food (Mery and Kawecki, 2004; Riffell et al., 2008; Gong, 2012). Because innate behavioral plasticity and learning both refer to labile traits, i.e., to phenotypes that can change multiple times during an individual’s lifetime (in opposition to nonlabile traits, or developmental plasticity), they should in principle both provide an advantage in within-generation varying environments (Gomulkiewicz and Kirkpatrick, 1992). Thus, more work is needed to disentangle the effects of environmental patterns on the evolution of the different forms of plasticity. In particular, we need another discriminating factor than environmental variability to understand the specific advantage of learning over innate behavioral plasticity. Such a distinction has not been made possible in previous theoretical work because in most models for the evolution of learning, learners are pitted against individuals that can only express one given genetically determined action (Boyd and Richerson, 1988; Stephens, 1991; Feldman et al., 1996; Kerr and Feldman, 2003; Wakano et al., 2004; Dunlap and Stephens, 2009). But this is not a very likely evolutionary transition. Learning is more likely to evolve on top of innate behavioral plasticity (Kerr, 2007) and it is indeed common to observe the coexistence within individuals of these two forms of plastic responses (Mery and Burns, 2010; Snell-Rood, 2013).

In this paper, our aim is to investigate the evolutionary transition from innate behavioral plasticity to learning, and show that learning is adaptive under conditions of environmental complexity. By environmental complexity, we mean the number of distinct challenges or stimuli that an individual encounters within its lifespan. These may be, for instance, an encounter with a predator or with a food item of some nutritional value. The reason why environmental complexity may select for learning can be verbally explained as follows. Because the range of challenges encountered during an individual’s lifepsan can be extremely large, and each of these situations generates a particular combination of sensory perceptions in the animal’s brain, it seems unlikely that an animal is capable of storing the interactions with all these challenges and the associated responses (even with the abstract representation provided by neural networks, Enquist and Ghirlanda, 2005).

The discrepancy between the complexity of the environment and the capacity of an individual’s memory to process information thus imposes a computational constraint on its decision system. Having a dynamic memory, which allows to forget obsolete stimulus-response associations and learn new ones may be useful to deal with environmental complexity, as it makes feasible to react to an arbitrarily large number of situations. The contribution of forgetting to the adaptive value of learning has already been investigated (Kraemer and Golding, 1997; Kerr and Feldman, 2003), but only in situations where forgetting allows one to face the same challenge at distinct instants and if the optimal behavior for that challenge has changed (i.e. environmental variability). We rather propose that forgetting contributes to the benefits provided by learning through the ability to encode different stimuli, because different stimuli may be encountered at distinct instants of time. This is consistent with the functioning of short-term memory: animate or inanimate features with which an animal interacts first enter the working memory and are transferred to the long-term memory only through a consolidation phase, which does not necessarily occur (Dudai, 2004; Shettleworth, 2009). When supplemented with forgetting, learning is thus likely to provide a powerful mean to cope with environmental complexity, because it can scatter complexity over time; only a small portion of the environment’s complexity is dealt with per unit time.

In the rest of the article, we formalize in an evolutionary model the above verbal argument that instrumental learning is adaptive under conditions of environmental complexity. In order to capture the limitations of an individual in terms of information processing, we assume that it is constrained by a maximum amount of memory. An evolving trait prescribes the allocation of this memory either to an innate memory or to a dynamic memory, which allows the individual to learn and forget associations between stimuli and actions. The environment consists of a finite (but possibly very large) number of challenges (or stimuli), each of which is characterized by its own optimal action(s). We will show that environmental complexity (operationalized as the number of potential stimuli in the environment) can generate a selection pressure in favor of a greater allocation of memory to learning.

## Model

### The individual and its environment

Consider an individual that interacts with its environment for successive discrete time steps. At each time step, the individual has to choose an action to respond to an environmentally determined challenge or stimulus, which is drawn from a set of *N*_s_ stimuli. Following previous formalizations (Feldman et al., 1996; Wakano et al., 2004), we assume that the chosen action is either the “correct” (or appropriate) response to the stimulus and gives payoff *π_C_*, or is a “wrong” response and gives payoff *π_W_*.

Because stimuli may depend on location, task to be performed, or time of the day, the individual is unlikely to meet all of them at once and we assume the following environmental process, where only a subset of the entire set of stimuli can be encountered per time step. Namely, any time step of an individual’s lifepsan consists of three events. (1) With probability *γ* a block of stimuli of size *N*_b_ ≤ *N*_s_ is randomly drawn from the set of environmental stimuli, while with probability 1 − *γ* the invidual faces the same block met at the previous time step. (2) A stimulus is uniformly drawn from the block (hence a given stimulus in the block is sampled with probability 1/*N*_s_). (3) The individual chooses an action in response to this stimulus.

An important property of this environmental process is that the stationary distribution of stimuli is uniform, so that every stimulus has probability 1/*N*_s_ of being encountered in the stationary state (see the Electronic supplementary file 1 (ESM1) for a proof and a detailed mathematical description of the environmental process). We take *N*_s_ to be a measure of *environmental complexity:* large values of *N*_s_ correspond to the case where an individual will encounter many different stimuli. However, the environment itself is not uniform and the parameter *N*_b_ (1 ≤ *N*_b_ *≤ N*_s_) captures *local complexity*: low values of *N*_b_ correspond to a low number of stimuli possibly experienced per interaction period, while high *N*_b_ corresponds to a locally complex environment. Finally, the parameter *γ* (0 ≤ *γ* ≤ 1) is the *environmental switching rate:* low values of *γ* correspond to small block turnover. In this environment, predictability, a feature that has been shown to critically affect the evolution of learning (Stephens, 1991), is captured by the interaction between *N*_b_ and *γ*. When *γ* is close to 1, there is high block turnover, so the environment is not very predictable for any block size *N*_b_. When *γ* is smaller, predictability depends on local complexity, *N*_b_. A large value of *N*_b_ means that a lot of stimuli are being encountered in a given block so the probability to encounter the same stimulus repeatedly is low, and hence the environment is less predictable. A small value of *N*_b_ corresponds to a more predictable environment where the individual only deals with a small number of stimuli during a period of interaction with a block.

In order to be able to store information about how to respond to stimuli, we assume that the individual has a memory that can store *m* associations between stimulus and action. These *m* memory “slots” could either be filled with fixed associations present at birth, which hold templates of stimuli together with the innate response to these stimuli, or with such associations that are learned during the individual’s lifespan. We denote by *g* the number of associations that are innately determined. Hence, if *g < m*, a part of the memory, *m − g* slots, is dynamic, and the individual can encode new stimuli encountered during its interactions with the environment. If *g* = 0, the individual is born with a “blank slate”, with absolutely no innate tendency to respond to environmental stimuli. Our goal is to understand the selection pressure on the evolving trait *g*, given a fixed memory capacity *m*, and how this depends on environmental complexity (*N*_s_), local complexity (*N*_b_), and switching rate (*γ*).

### Fitness

In order to evaluate the selection pressure on g, we need a measure of expected payoff (or fitness) accruing to an individual expressing this trait value. To obtain this, we note that an encountered stimulus at a given time step can be in three possible states with regard to the individual’s memory. First, the stimulus can be innately encoded, in which case we denote by *π_I_* the average payoff obtained from the response to it. Second, the stimulus encountered can be present in the dynamic memory, in which case the response results in average payoff *π_L_.* Third, the stimulus may not be present at all in the memory of the individual (it is “unknown”), in which case the individual’s response results in average payoff *π_U_*.

Because our main interest is in understanding the environmental conditions that favor learning, we assume (conservatively) that expressing a genetically determined action always leads to the “correct” payoff: *π_I_ = π_C_*. When an individual encounters a stimulus that is not in its memory, we assume it samples an action at random. The expected payoff obtained by choosing an action randomly is denoted *π_U_*, and is assumed to satisfy *π_W_ ≤ π_U_ ≤ π_C_*. Finally, we assume that *π_L_*, the payoff for learned responses to stimuli, is a constant satisfying *π_U_ ≤ π_L_ ≤ π_C_*, so that an action for a stimulus present in dynamic memory leads to a higher payoff than if it was tried out randomly, because learning allows to sample the environment. A distinctive simplifying feature of our model, which gives analytical traction, is that we do not model explicitly the learning dynamics of such association between actions and stimuli (for instance by way of reinforcement learning). But by enforcing *π_U_ ≤ π_L_ ≤ π_C_*, we implicitly capture any learning mechanism, from a very crude one where essentially no information is gathered if *π_L_ ≈ π_U_* to a very sophisticated one if *π_L_ ≈ π_C_*.

Owing to the assumption that the stimuli are met by the individual in a stationary uniform distribution, the probability *P_I_*(*g*) that a currently encountered stimulus is innately encoded is independent of time; namely *P_I_*(*g*) = *g/N*_s_. Assuming that the individual interacts a very long time with its environment, we have that the asymptotic probability, *P_L_*(*g)* that an encountered stimulus is in the dynamic memory is also independent of the time (this assumption can indeed be justified for the environmental process we consider in this paper, see the ESM1). With this, we can then write the average payoff to an individual with genetic memory of size *g* as

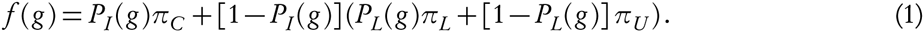

This equation captures the trade-off faced by the individual: should it allocate memory to innate responses and respond optimally to only a limited number of stimuli (first term of eq. 1), or should it allocate memory to learning, and potentially learn to respond to many stimuli (second term of eq. 1)? Importantly, if the stimulus recall probability, *P_L_*(*g)*, is a constant, independent of *g*, then the optimal number of innate memory slots, *g*^*^, which maximizes payoff, is just *g*^*^ = *m*. Hence learning does not evolve in this case (the same holds if *P_L_*(*g)* is increasing in *g*). This function thus requires that *P_L_*(*g)* is decreasing in *g*, at least on some subset of [0, *m*], for learning to evolve. But the exact form of *P_L_*(*g)* will depend on how memory works, i.e. for how long a stimulus is stored in memory before it is forgotten.

## Memory

We endorse a simplified implementation of memory that is based on the functioning of the short-term memory in humans and animals (Baddeley, 2003). Namely, we assume that, when the individual has a dynamic memory (*g* < *m*) and meets an unknown stimulus, it always wants to store it. Since the dynamic memory is initially empty, the first *m − g* encounters with non-innately encoded stimuli will simply result in the stimuli taking free slots until the dynamic memory is full. For subsequent decision steps, new stimuli will have to replace other ones in the dynamic memory. This is done via a replacement rule. We use the following replacement rule, which is taken from Kerr and Feldman (2003). A stimulus has a lifespan in memory of *m − g* time steps (i.e. the size of the dynamic memory), starting from the last encounter with the stimulus. This means that if a stimulus is not met more than once in *m − g* steps, it is forgotten. Otherwise, the stimulus stays in memory. With this rule, the dynamic memory will never contain more than *m − g* stimuli.

Importantly, we assume that when a stimulus is replaced in memory, all the associated information is lost (note that this is another very conservative assumption, because this means that we ignore the potential benefits of long-term memory). If this stimulus is encountered later, then the individual will have to re-learn to interact with it (if the individual has the capacity to do so, i.e. if *g < m*). With this replacement rule, we can now evaluate *P_L_*(*g)* explicitly. We are then able to ascertain the evolutionarily stable value of *g* by taking the expected payoff (eq. 1) as our measure of fitness (Parker and Maynard-Smith, 1990).

## Results

### Stimulus recall probability

In the ESM1, we derive an expression for the stimulus recall probability, *P_L_*(*g)* (eq. S7). It turns out that this expression is cumbersome (see ESM1, section “Stimulus recall probability”, to observe the full expression), but in Fig. 1, we plot *P_L_*(*g)*, which shows that it is decreasing in *g*, and when *g* = *m* we have *P_L_*(*m)* = 0. This decreasing pattern obtains because having a higher *g* means having less slots for the dynamic memory, which in turn implies that an individual will recall less steps of interaction with a given stimulus. The stimulus recall probability depends not only on the memory characteristics of the individual (*m* and *g*), but also on the three key environmental parameters of the model: environmental complexity (*N*_s_), local complexity (*N*_b_), and switching rate (*γ*).

**Figure 1.**
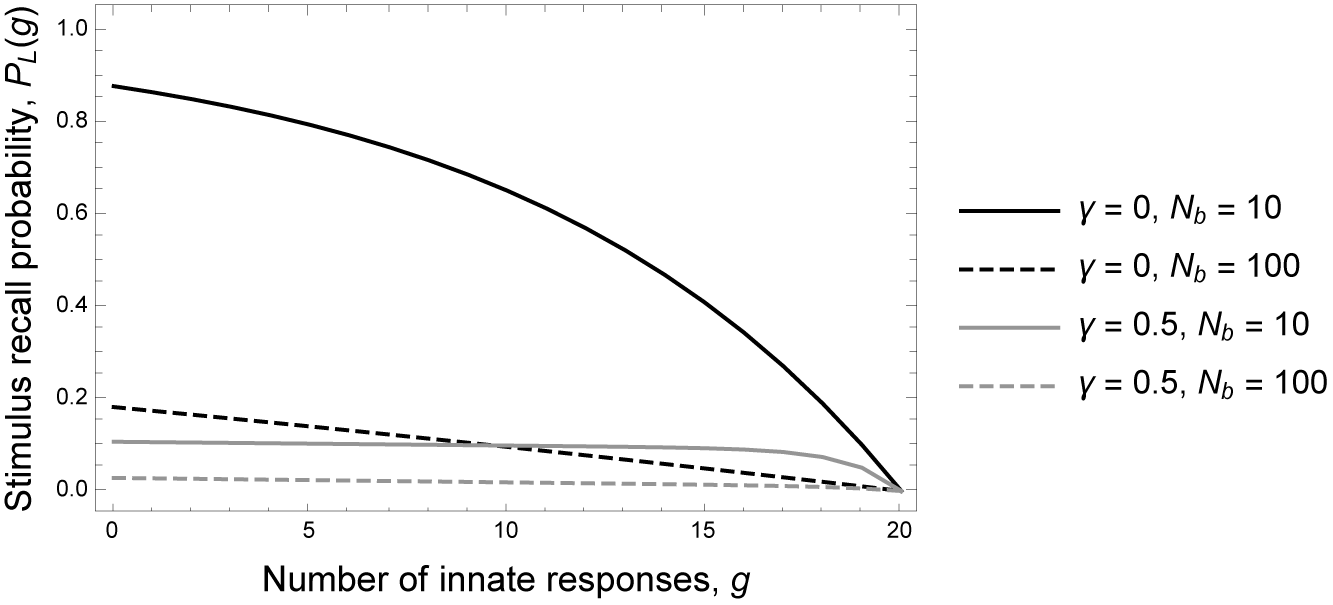
Stimulus recall probability, *P_L_*(*g)*, as a function of number of innate responses, g, and environmental parameters. Parameter values: *m* = 20, *N*_s_ = 1000.

As can be seen from Fig. 1, *P_L_*(*g)* is decreasing in *γ*, which stems from the fact that when the block of stimuli changes more frequently, the probability to encounter multiple times the same given stimulus decreases (lower predictability). We also have that *P_L_*(*g)* is smaller in locally complex than in locally simple environments, because greater local complexity corresponds to more stimuli in a block. Finally, *P_L_*(*g)* is slowly decreasing with increasing *N*_s_ and eventually stabilizes for large *N*_s_. This is mainly due to the fact that when the environment is complex, there is a very small probability that a given stimulus is in two different blocks of stimuli, so an individual will recall mainly interactions with stimuli within blocks, not across blocks.

The stimulus recall probability thus has two main properties, one related to the memory of the individual, and the other one related to the environment: it is an increasing function of the allocation of memory to learning, and also generally increases in the predictability of the environment.

### Invasion of learners

When will learning be initially favored by selection? To answer this question, we consider a monomorphic population of “innates” with *g* = *m* and ask when they will be invaded by mutant learners with only one memory slot allocated to the dynamic memory (*g* = *m* − 1); that is, when *f* (*m) < f* (*m −* 1) is satisfied. This occurs when

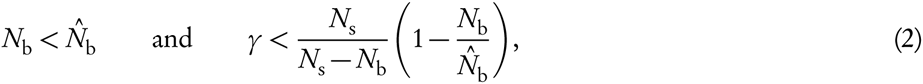

where

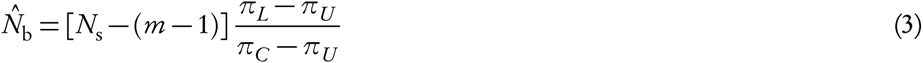

(see the ESM1 for a proof). This can be thought as the total gain from having a dynamic memory for an individual characterized by only one memory slot dedicated to learning (*g* = *m* − 1). Indeed, *N*_s_ − (*m* − 1) is the total number of environmental stimuli that are not innately recognized by the individual (and thus that can be learned about). The ratio (*π_L_ − π_U_)/*(*π_C_ − π_U_*) varies between 0 and 1 (because *π_U_ ≤ π_L_ ≤ π_C_*) and represents a normalized gain from learning to interact with an unknown stimulus. Making the environment more complex (increasing *N*_s_) widens the range of parameters where learning invades, because this makes larger the threshold 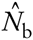 below which learning evolves (Fig. 2). It is important to emphasize that even when the payoff due to learning, *π_L_,* is only slightly higher than the random payoff, *π_U_*, environmental complexity still has a positive effect on the evolution of learning; one just needs to make the environment complex enough for ineq. 2 to be satisfied. Hence, the model captures well the evolutionary transition from optimal innate behavioral plasticity to an imperfect learning system, which is in theory the possible first state of a learning ability.

**Figure 2.**
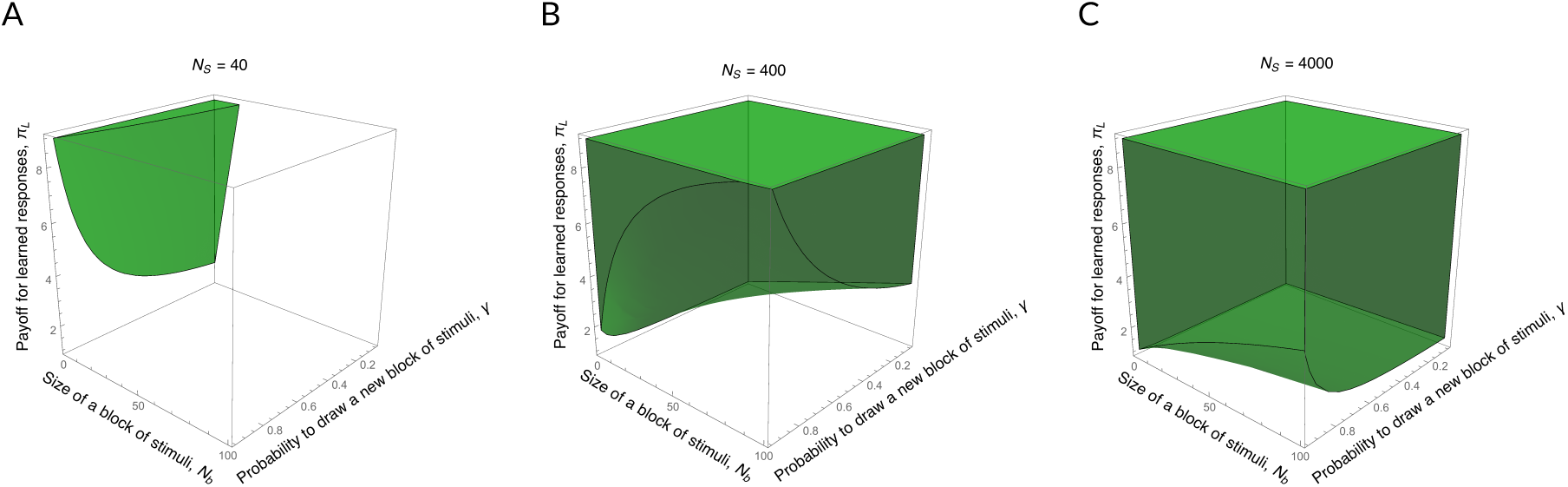
Increasing volumes where learning invades as a function of environmental complexity. In each subfigure (A,B,C), the green volume corresponds to the region of the (*γ*, *N*_b_, *π*_L_)-parameter space where a mutant learner invades a monomorphic population of individuals relying on innate behavioral plasticity (i.e. ineq. 2 is satisfied). Parameter values: *m* = 20, *π_C_* = 10, *π_U_* = 1. A: *N*_s_ = 40. B: *N*_s_ = 400. C: *N*_s_ = 4000.

Ineq. 2 also shows that the environment should be predictable enough for learning to be favored by selection. In terms of our model parameters, this translates as a maximal amount of environmental switching rate (*γ* small enough), and as a maximal local complexity (*N*_b_ smaller enough than *N*_s_). Indeed, if *N*_b_ = *N*_s_, there is only one block of stimuli of size *N*_s_, and the individual must cope with the total amount of complexity at once; a learner with a limited memory size cannot cope with such a task. Also, *γ* should be small enough so that the average period of interaction with a block, 1/*γ* is large enough compared to the local complexity, *N*_b_. The threshold value of *γ* in ineq. 2 is decreasing when *N*_b_ increases, which indicates that locally complex environments require longer interaction periods with blocks. This will allow a learner to interact many times with the same stimulus and learn the best response to it.

## Optimal memory allocation

We now turn to investigate numerically the optimal value *g*^*^ that maximizes fitness (eq. 1); that is, the evolutionarily stable allocation of memory to innate behavioral plasticity. First and foremost, we find that increasing the complexity of the environment increases the optimal size of the dynamic memory: *g*^*^ is decreasing with increasing *N*_s_. This is because in complex environments, where *N*_s_ is much larger than the memory size *m*, allocating one more slot to the innate memory has only a small effect on fitness (first term of eq. 1); by contrast, allocating this slot to the dynamic memory always results in an increase of the stimulus recall probability, even for large *N*_s_ (second term of eq. 1 and Fig. 1). Likewise, increasing the efficiency of the learning system, *π_L_,* also decreases the value of *g*^*^ (Fig. 3A) because increasing *π_L_* means having a higher benefit of learning (i.e. this increases the value of the second term of eq. 1) for an individual with a given *g*.

**Figure 3.**
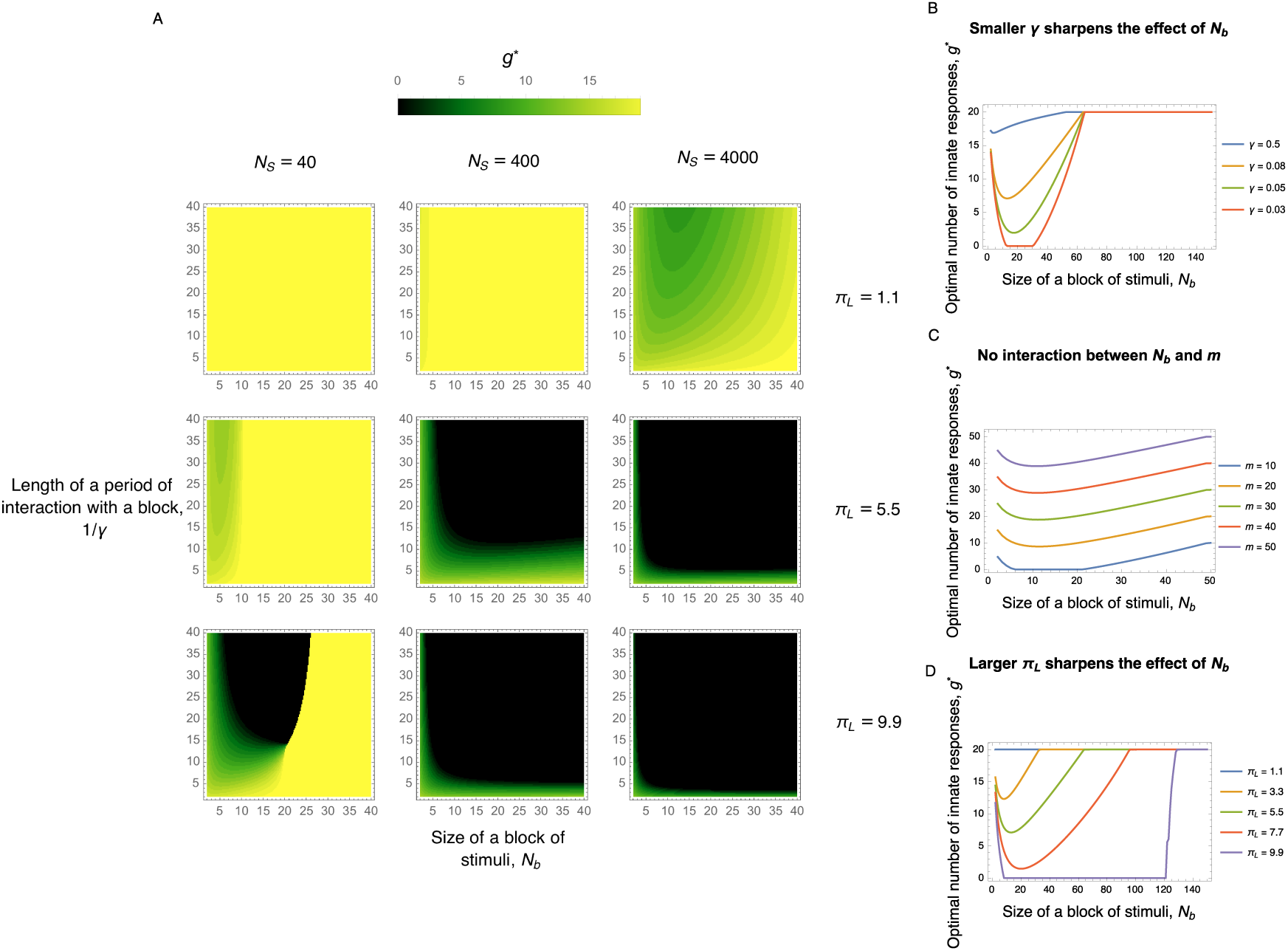
Optimal number of innate responses, *g*^*^, that maximizes eq. 1, as a function of environmental parameters and efficiency of learning mechanism. A: Concomitant effects of environmental complexity (*N*_s_, columns), efficiency of learning mechanism (*π_L_*, rows), local complexity (*N*_b_, *x*-axis), and environmental switching rate (*γ*, *y*-axis) (parameter values: *m* = 20, *π_C_* = 10, *π_U_* = 1). B: Interaction between *N*_b_ and *γ* (parameter values: same as in A, plus *N*_s_ = 150, *π_L_* = 5.5). C: Interaction between *N*_b_ and *m* (parameter values: same as in A, plus *N*_s_ = *m* + 100, *π_L_* = 5.5, *γ* = 1/12). D: Interaction between *N*_b_ and *π_L_* (parameter values: same as in A, plus *m* = 20, *N*_s_ = 150, *γ* = 1/12).

The environmental switching rate, *γ*, has a simple effect on optimal memory allocation. In agreement with the invasion condition (ineq. 2) and the above fact that the probability to recall a stimulus is decreasing in *γ*, the optimal number of innate responses, *g*^*^, is increasing in *γ*. This makes sense since in an environment changing more frequently, it is less beneficial to recall events farther in the past. However, there is a threshold effect when we have no environmental change at all (*γ* = 0), and the individual is faced with only a random sample from the environment for its entire lifespan. This leads to a null model where the environment is totally random and learning is helpless.

Local complexity, *N*_b_, has a non-monotonic effect on *g*^*^ (see in particular Fig. 3B,C,D). For locally complex environments, *g*^*^ is increasing, while for locally simple environments it is decreasing. Hence, the maximum allocation of memory to learning occurs at moderate levels of local complexity. This pattern is explained in terms of the marginal gains of allocating memory slots to either part of the memory. When *N*_b_ is high, the gains from allocating a memory slot to the innate memory are higher than the gains from allocating it to the dynamic memory because there is only a small probability to recall interactions with stimuli (low predictability). When *N*_b_ is low, the converse is true: it is more beneficial to allocate a slot to the dynamic memory because the stimulus recall probability is high (high predictability).

We also looked at various two-way interactions between parameters. First, it is interesting to investigate the interaction between the switching rate and local complexity (Fig. 3B) because together they determine the predictability of the environment. We find that the environmental switching rate sharpens the non-monotonic effect of *N*_b_ described above. This is because in fast changing environments, there is very low predictability irrespective of local complexity, since there is only a small probability to encounter twice the same stimulus (in this case, *P_L_*(*g)* is flat, Fig. 1). In slowly changing environments where interaction periods are longer, the stimulus recall probability is now highly dependent on *N*_b_ and *g*. Hence for slowly changing environments (low *γ*), high *N*_b_ corresponds to low predictability and low *N*_b_ to high predictability, and we recover the non-monotonic effect of *N*_b_ found above.

Strikingly, there is no interaction effect between local complexity and memory size on *g*^*^, provided *N*_s_ − m is kept constant (Fig. 3C). This is intriguing because one would think that if the environment is locally complex (*N*_b_ large), then there is a very small probability to interact many times with a given stimulus in a short period of time, thereby rendering learning more difficult for an individual with a given memory *m*. However, making *N*_b_ large means making *N*_s_ at least as large, so the interaction between *N*_b_ and m is already captured by the main effect of *N*_s_ − *m*, which explains why there is no interaction effect between *N*_b_ and *N*_s_ − *m* on *g*^*^. The payoff for learned responses to stimuli, *π_L_,* finally, has the role of making the effect of the other parameters more abrupt. For instance, when *π_L_* increases, we observe that below the threshold value *N*_b_ of block size favoring the invasion of learners (ineq. 2) all memory slots are dedicated to learning (*g*^*^ = 0, Fig. 3D).

## Discussion

In this study, we investigated the evolutionary transition from innate behavioral plasticity to learning, and showed with an analytical model that environmental complexity (operationalized as the number of stimuli in the environment, *N*_s_) favors the evolution of learning. Because we considered an environment that is constant across generations, yet where learning can invade, our results demonstrate that between-generation environmental variability is not necessary for learning to evolve. Further, we found broad conditions where learning coexists with innate behavioral plasticity (i.e. 0 *< g*^*^ *< m*).

### Summary of results

Our results are twofold. First, we provide conditions for learners to invade a population of individuals relying on innate behavioral plasticity. We find that increasing the complexity of the environment widens the range of conditions under which learners can invade. This is due to the fact that with innate behavioral plasticity in a complex environment, an individual cannot form new stimulus-response associations during its lifespan, and thus can respond to only a limited number of stimuli. We also find that environmental complexity is not a sufficient condition for learning to evolve. Namely, we confirm previous results showing that the environment needs to display predictability within an individual’s lifespan (Stephens, 1991; Dunlap and Stephens, 2009, 2014). In locally complex environments, where an individual interacts with blocks consisting of many stimuli at the same time, predictability is very low, i.e. there is a small probability to encounter the same stimulus in a short period of time. In this case, learning cannot invade because the limited memory of the individual is unable to deal with the entire environmental complexity at once.

Our second types of results are related to the optimal allocation of memory between innateness and learning. In our model, this optimal allocation is determined by the trade-off between using memory to respond optimally and innately to only a few stimuli, versus using this memory to learn to respond sub-optimally to potentially many stimuli. As could be anticipated from the invasion results, we find that the optimal allocation of memory to learning increases with the complexity and the predictability of the environment. Moreover, we find that the maximum allocation to learning occurs in moderately predictable environments. These are the environments where block turnover is small (i.e. small switching rate) but where local complexity (or block size) is intermediate. In these cases, we even find conditions where it is optimal that individuals are born with a “blank slate” (all memory slots allocated to the dynamic memory). Overall, the results show that learning supplemented with forgetting represents an efficient way to deal with environments that are complex on the global scale but are relatively simple on a local scale. But how can we measure complexity in the real world, and what phenotypes are affected by it?

### Empirical predictions

The expression “environmental complexity” used in this article refers to the number of fitness-relevant stimuli a given individual is likely to encounter and distinguish in the course of its lifespan. There are at least three measurable ecological and psychological factors that directly or indirectly influence complexity. First, the complexity of an organism’s environment should be positively correlated with the habitat range of that organism. Individuals from species exploring vast areas in order to forage, migrate, or reproduce should encounter more types of biotic and abiotic stimuli than individuals from other species. Second, the level of detail that an organism’s sensory system can perceive is also likely to allow an individual to distinguish between many stimuli (typically, species with a visual ability that are color-blind “miss” one dimension of the world’s complexity). Third, lifespan is a factor that will affect the number distinct stimuli or challenges encountered by a given individual: species with longer lifespan should be exposed to a greater variety of stimuli.

Our results thus predict that these factors should be positively correlated with learning ability. Mainly, we expect that species scoring very low on the three dimensions of complexity highlighted above should be those species that have a scant ability to learn, and rely on simpler forms of plasticity (that we termed “innate behavioral plasticity”). More marginally, our results suggest that the capacity (or size) of individuals’ working memory is likely to increase with complexity. This number corresponds to the number of stimuli (or chunks of information) an individual can hold in short-term memory for further processing and use (Carruthers, 2013).

### Model realism

Our model is obviously a simplification, but we argue that the environment we considered is representative of those faced by many animals. To give a concrete example of the range of settings where our model applies, we can take daily routines (Houston and McNamara, 1999). In each part of its routine, an animal interacts with a given subset of stimuli that are present at a given location and time, because of statistical regularities in the environment. For example, when an individual visits a particular food patch in the course of foraging, it may encounter different types of food items, but also individuals from other species that have overlapping diet, as well as predators awaiting for their preys. All of these constitute the block of stimuli met by the individual on this particular food patch; upon visiting other food patches and performing other tasks, the individual will encounter other blocks of stimuli (that may or may not contain the stimuli previously encountered).

While our model is general enough to capture features of real environments, our implementation of memory is relatively specific. In order to ground our results on the most possible conservative assumptions, we made two notable simplifications. First, we focused on instrumental rather than associative learning. In artificial selection experiments where the evolution of learning was shown to be favored by between-generation environmental variability (Mery and Kawecki, 2004; Dunlap and Stephens, 2009, 2014), it was associative rather than instrumental learning that was considered. It is likely that the conditions favoring associative learning are different than the ones favoring instrumental learning. However, in most models of the evolution of learning, the modeling approach is so abstract that it may encompass both forms of learning (Boyd and Richerson, 1988; Stephens, 1991; Feldman et al., 1996; Wakano et al., 2004). These mechanistic considerations need further evolutionary investigation and clarification.

Our second simplification is that we considered only features of the short-term (or working) memory (Shettleworth, 2009; Banai et al., 2010). In contrast with long-term memory, the events stored in working memory follow a dynamic process such that they enter memory when an individual starts interacting with a given stimulus, but such interactions are replaced by other ones as the individual interacts with different stimuli. Indeed, animals and humans can hold only a given, small number of items or stimuli in working memory (Miller, 1956; Dudai, 2004). This is captured by the dynamic part of the memory in our model, where stimulus-action associations are totally removed from memory once they are replaced by other ones.

The “innate” part of the memory, on the other hand, captures many of the examples in nature showing that animals tend to have innate, hard-wired responses to stimuli (Mery and Kawecki, 2004; Riffell et al., 2008; Gong, 2012). For instance, certain ants have innate templates of enemies in memory (Dorosheva et al., 2011) and human infants innately distinguish between face-like stimuli and other stimuli, indicating that the neuronal networks responsible for visual perception have a particular innate wiring structure (Slater and Kirby, 1998; see also Perin et al., 2011 on a generalization of this idea to the innate structure of the whole neocortex of mice).

In conclusion, this study provides a new perspective on the role of environmental complexity in the evolution of learning mechanisms (Jones and Blackwell, 2011; Fawcett et al., 2014). This work also provides a clarification on the role of predictability in this process, and show that these elements generate environmental dynamics within a individual’s lifepsan that are useful to take into account in order to understand the evolution of learning and memory.

## Supporting Information for “Environmental complexity favors the evolution of learning” by Slimane Dridi and Laurent Lehmann

Here, we derive an expression for the probability to recall a stimulus *P_L_*(*g)*. At each time step *t* (*t* = 1,2,…) of the environmental process, a stimulus *s_t_* is drawn from the set of environmental stimuli, *S*, according to the procedure described in the main text (i.e. *s_t_* is a random variable). Now, given our implementation of memory, where a given stimulus stays in dynamic memory *m − g* time steps, *P_L_*(*g)* is the probability that, conditional on being met once, this stimulus is met a second time in a period of length less or equal than *m − g* time steps. Let us denote by *R_s_* ∈ {1,2,…} the number of time steps occurring between two encounters with a given stimulus (this is a random variable that is independent of time at stationarity). Then

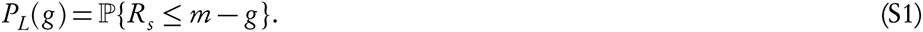

We will compute this expression by letting *R_s_* be the return (or recurrence) time of a backward Markov chain (Grimmett and Stirzaker, 2001) that can be constructed from our assumptions on the environment.

To that end, let 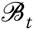 ⊂ *S* be the block of stimuli 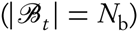 encountered at time *t*,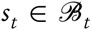 the stimulus encountered at *t*, and *S_f_* ∈ *S* a given focal stimulus. We can then define the three mutually exclusive events

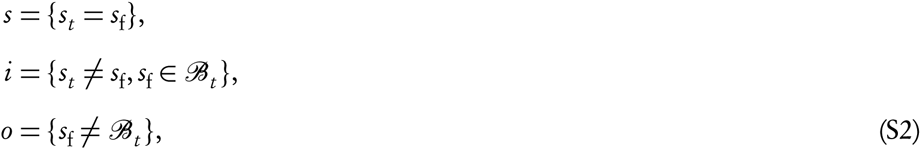

where *s* is the event that the stimulus encountered at time *t* is the focal one, *i* is the event that the focal stimulus is in the current block but is not the currently encountered stimulus, and *o* is the event that the focal stimulus is not in the current block.

We can now define a Markov chain on these three states: *s*, *i*, and *o* and compute from it the recurrence time to *s*. From our assumptions, the forward transition probabilities *ρ_jk_* of this chain are

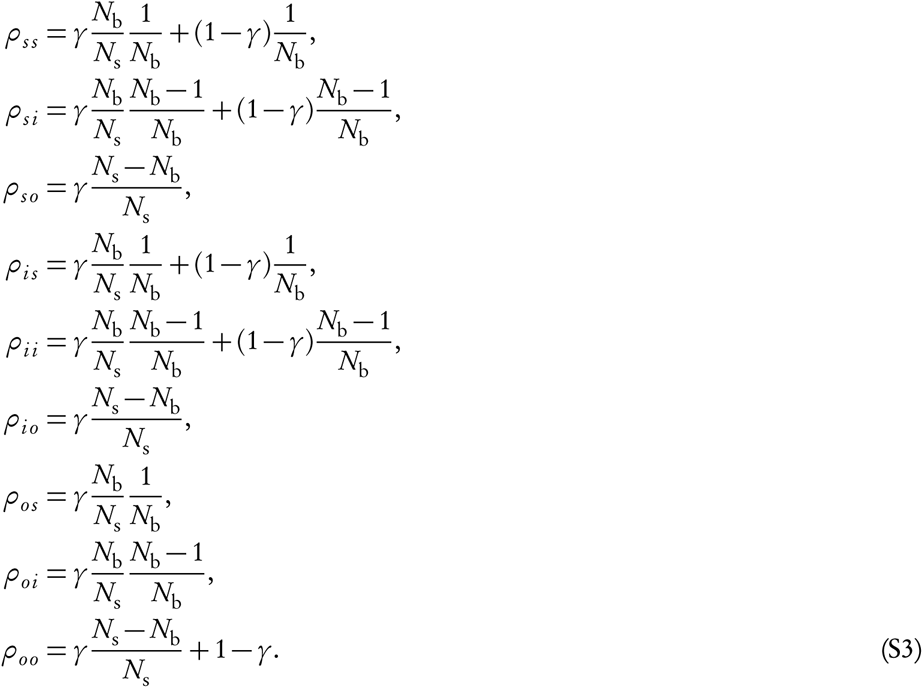

For instance, the probability *ρ_ss_* to move from state *s* to itself takes this form because a new block is drawn with probability *γ*, in which case the focal stimulus *s*_f_ makes part of the new block with probability *N*_b_/*N*_s_, and is drawn from within the block with probability 1/*N*_b_. If one does not change block, which happens with probability 1 − *γ*, the probability to draw *s*_f_ from the current block is 1/*N*_b_, because *s*_f_ already makes part of the current block.

From the above transition probabilities, we can define the backward transition probabilities *pkj*, and owing to our stationarity assumption, this is given by

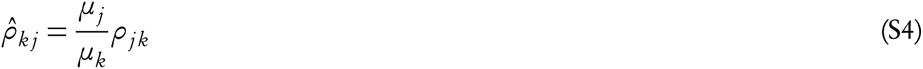

(see e.g. Theorem 1.9.1 of Norris, 1998), which defines a backward Markov chain. Since the stationary probabilities are given by *μ_s_* = 1/*N*_s_, *μ_i_* = (*N*_b_ − 1)/*N*_s_, and *μ_o_* = (*N*_s_ − *N*_b_)/*N*_s_, we find using eq. S3 that

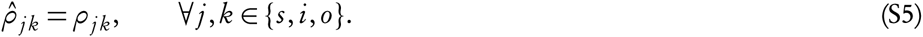

Now, we can compute the distribution of return times *R_s_*. To do so, denote 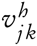 the probability that, starting from state *j*, the first visit to state *k* occurs *h* steps in the past. With this, we have that 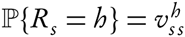. In order to find 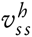, we note that the probabilities of first visit obey the recursions

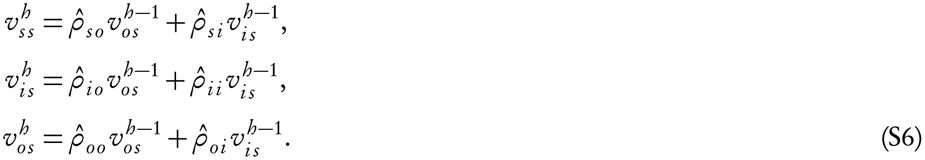

Solving this linear system of difference equations provides the probabilities of first visit on the left-hand side, including 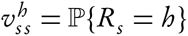 (we display this long expression at the end of this Supplementary file, section “Distribution of return times”). The probability that *R_s_* ≤ *m − g* (*m − g* > 0) can then be computed by summing all the possible cases up to *m* − *g*, namely

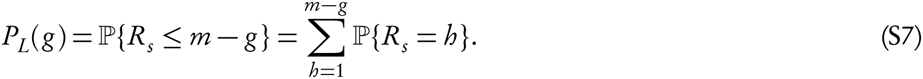

It turns out that, substituting the explicit expression of 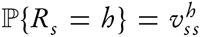, we obtain a closed-form expression, but that is unfortunately too long to provide direct insight (see the section “Stimulus recall probability” at the end of this Supplementary material).

### Invasion and stimulus recall probabilities

In this section, we show how to derive the invasion condition in ineq. 2. Applying the derived probability found above (eq. S7 and section “Stimulus recall proabability” below), we have that *P_L_*(*m*) = 0 and

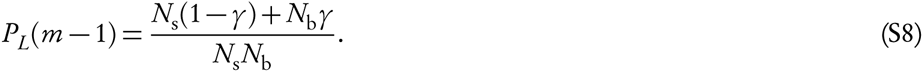

Substituting these in *f* (*m* − 1) > *f* (*m*) gives ineq. 2 of the main text.

### Distribution of return times (eq. S6)

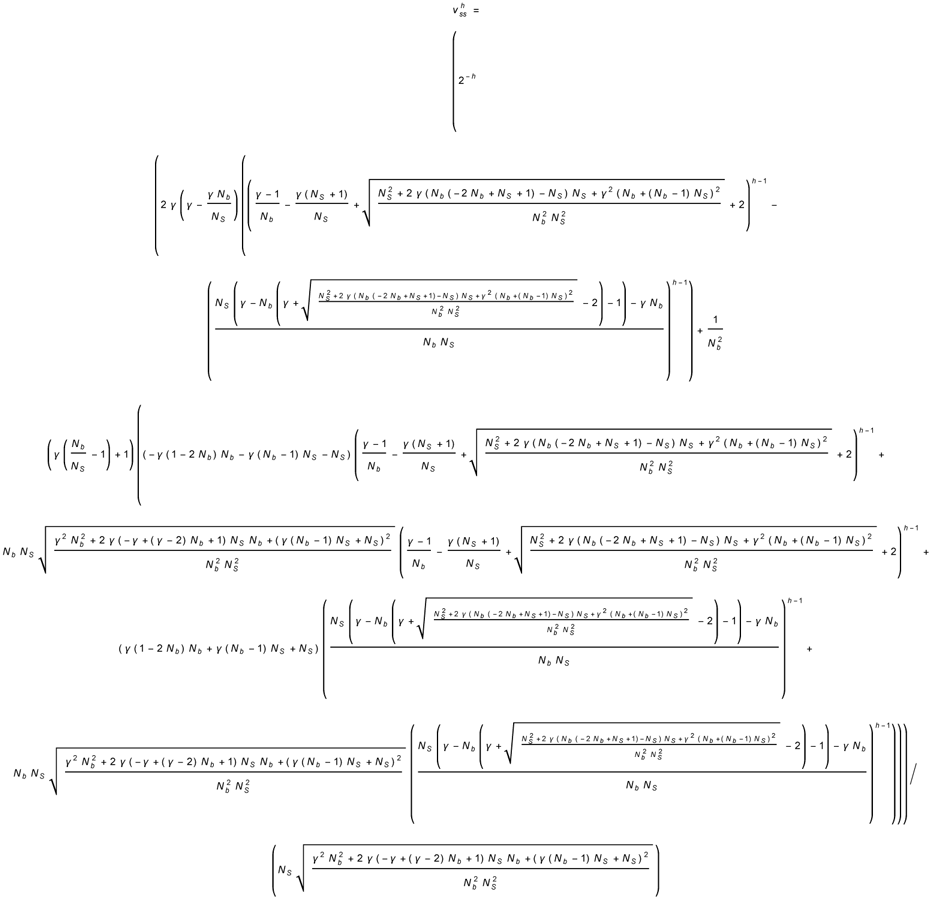

### Stimulus recall probability (eq. S7)

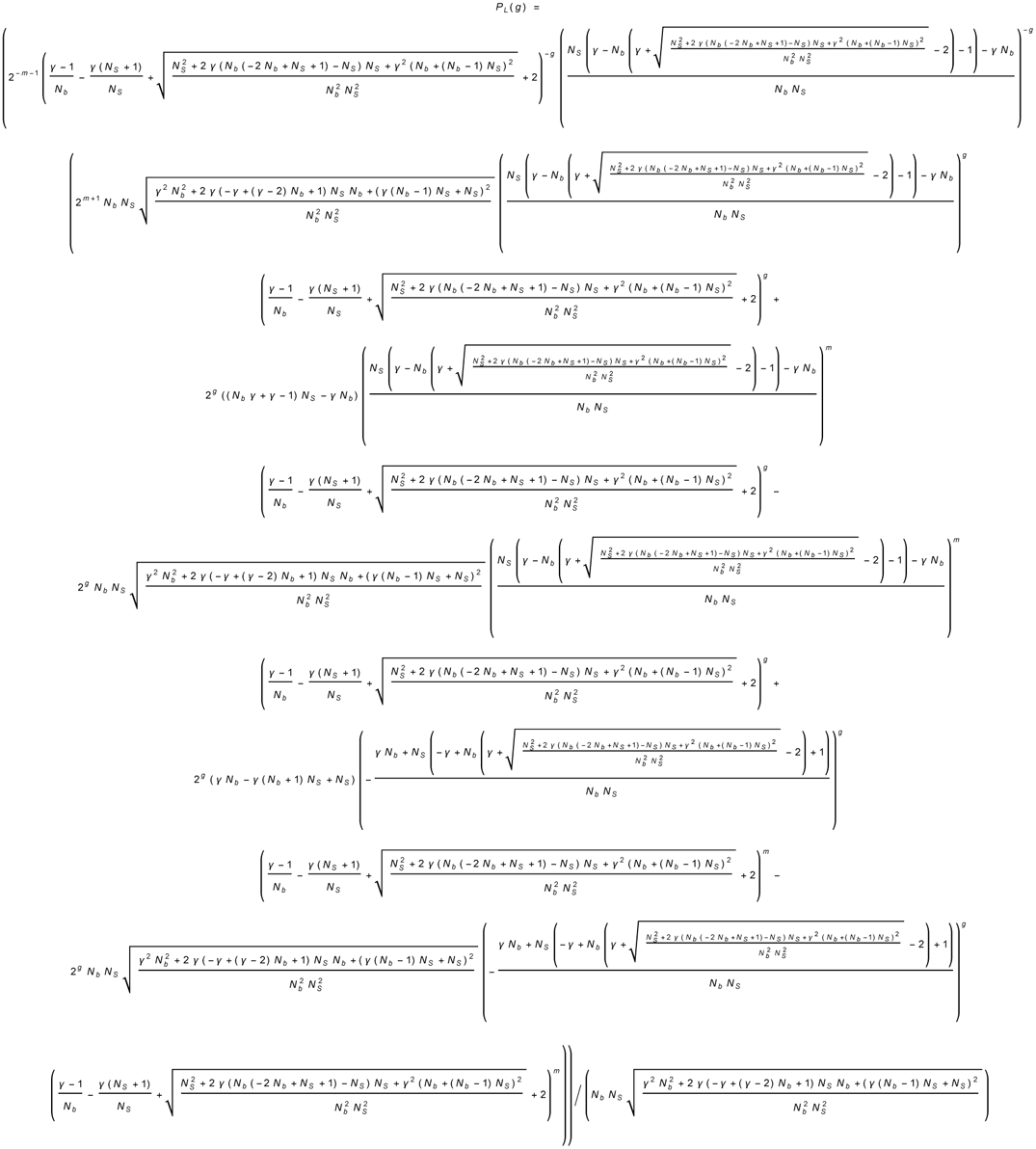

